# A mouse model of Bardet-Biedl Syndrome has impaired fear memory, which is rescued by lithium treatment

**DOI:** 10.1101/2020.10.06.322883

**Authors:** Thomas K. Pak, Calvin S. Carter, Qihong Zhang, Sunny C. Huang, Charles Searby, Ying Hsu, Rebecca Taugher, Tim Vogel, Christopher C. Cychosz, Rachel Genova, Nina Moreira, Hanna Stevens, John Wemmie, Andrew A. Pieper, Kai Wang, Val C. Sheffield

## Abstract

Primary cilia are microtubule-based organelles present on most cells that regulate many physiological processes, ranging from maintaining energy homeostasis to renal function. However, the role of these structures in the regulation of behavior remains unknown. To study the role of cilia in behavior, we employ mouse models of the human ciliopathy, Bardet-Biedl Syndrome (BBS). Here, we demonstrate that BBS mice have significant impairments in context fear conditioning, a form of associative learning. Moreover, we show that postnatal deletion of BBS gene function, as well as congenital deletion, specifically in the forebrain, impairs context fear conditioning. Analyses indicated that these behavioral impairments are not the result of impaired hippocampal long-term potentiation. However, our results indicate that these behavioral impairments are linked to impaired hippocampal neurogenesis. Two-week treatment with lithium chloride partially restores the proliferation of hippocampal neurons which leads to a rescue of context fear conditioning. Overall, our results identify a novel role of cilia genes in hippocampal neurogenesis and long-term context fear conditioning.

**Author summary:** The primary cilium is a microtubule-based membranous projection on the cell that is involved in multiple physiological functions. Patients who have cilia dysfunction commonly have intellectual disability. However, it is not known how cilia affect learning and memory. Studying mouse models of a cilia-based intellectual disability can provide insight into learning and memory. One such cilia-based intellectual disability is Bardet-Biedl Syndrome (BBS), which is caused by homozygous and compound heterozygous mutations of BBS genes. We found that a mouse model of BBS (*Bbs1*^*M390R/M390R*^ mice) has learning and memory defects. In addition, we found that other mouse models of BBS have similar learning and memory defects. These BBS mouse models have difficulty associating an environment with an aversive stimulus, a task designed to test context fear memory. This type of memory involves the hippocampus. We found that *Bbs1*^*M390R/M390R*^ mice have decreased cell production in the hippocampus. Treating *Bbs1*^*M390R/M390R*^ mice with a compound (lithium) that increases cell production in the hippocampus improved the learning and memory deficits. Our results demonstrate a potential role for cilia in learning and memory, and indicate that lithium is a potential treatment, requiring further study, for the intellectual disability phenotype of BBS.

## Introduction

Intellectual disability (ID) is one of the most common neurodevelopmental disorders, affecting 1% of the global population [1, 2]. Clinically, ID is characterized by a deficit in intellectual functioning and adaptive functioning [3]. There are limited pharmacological interventions for ID, partially due to the heterogeneous nature of ID, and a poor understanding of ID which can be attributed to a lack of animal models of ID [4, 5]. There is an urgent need to develop animal models to improve our understanding of the pathophysiological mechanisms underlying this pervasive condition.

Patients with abnormal cilia, i.e. ciliopathies, frequently present with ID, suggesting that cilia play an important role in learning and memory, yet the mechanisms underlying the phenotype remain unknown [6]. Primary cilia are microtubule-based structures that extend from the surface of nearly all cells in the body, including neurons. Cilia play a role in maintaining energy homeostasis and facilitating physiological responses to sensory stimuli [7]. There are robust mouse models of ciliopathies that recapitulate the primary features of the human ciliopathies, which allow us to study the role of cilia in learning and memory. To this end, we employed mouse models of the human ciliopathy, Bardet-Biedl Syndrome (BBS), which presents clinically with intellectual disability [8] in order to investigate the role of cilia in learning and memory.

BBS is a genetically heterogenous autosomal recessive ciliopathy with 22 known causative genes [9]. Clinical features of BBS include rod-cone dystrophy progressing to blindness, postaxial polydactyly, obesity, renal anomalies, and intellectual disability [10]. BBS proteins are involved in ciliary function. Eight BBS genes, specifically *BBS1, BBS2, BBS4, BBS5, BBS7, BBS8, BBS9*, and *BBS18* (*BBIP1*), encode the components of the BBSome [11, 12], an octameric protein complex. The BBSome regulates ciliary trafficking of G-Protein Coupled Receptors (GPCR) including SMO[13], NPY2R [14], MCHR1 and SSTR3[15], and D1R [16]), as well as non-GPCRs (TRKB[17]). Three non-BBSome BBS proteins (BBS6, BBS10, and BBS12) form a complex that mediates the assembly of the BBSome [18]. BBS3 is a GTPase that is also involved in ciliary receptor trafficking[19].

We have developed multiple mouse models of BBS [20]. Unlike some other ciliopathy mouse models, BBS models are viable and clinically relevant because they use mutations found in human patients [21–24]. We focused on the use of *Bbs1*^M390R/M390R^ mice, harboring the most common human BBS mutation, as it recapitulates many of BBS phenotypes present in patients, including obesity, retinopathy, and decreased hippocampal volume[24, 25]. Despite the phenotypic association between decreased hippocampal volume in patients and the known role of the hippocampus in learning and memory, the role of BBS in learning and memory is not well studied. Here, we investigate the role of these cilia genes in learning and memory using a fear conditioning paradigm.

Fear conditioning evaluates associative learning and involves pairing a neutral stimulus [conditioned stimulus (CS)], to an aversive stimulus [unconditioned stimulus (US)]. Fear conditioning is commonly used to understand the neurobiological mechanisms of ID as well as fear learning and memory in mice due to several advantages [26–29]. First, fear conditioning paradigms provide distinct insights into the neural correlates of learning and memory, for example, context or cue-dependent conditioning, which require contributions from different brain regions [30]. Second, the pairing of CS to US consistently elicits a measurable set of physiological and behavioral responses [31]. Third, fear conditioning allows for the delineation between short-term and long-term memory performance, depending on the time duration from training to testing. To assess short-term context fear conditioning, a one-hour interval between training and testing is utilized. To evaluate long-term context fear conditioning, an interval ≥ 24 hours is utilized [32–34]. Finally, fear conditioning is a form of passive learning, making it an accessible behavioral test for mice with motor deficits [35].

Here, we report that *Bbs1*^M390R/M390R^ mice have impaired long-term context fear conditioning, but normal short-term context memory. In addition, we show that multiple BBS mouse models have impaired long-term context fear conditioning, including a mouse model with preferential deletion of *Bbs1* in the forebrain. We also show a novel role for BBS genes in neural proliferation and neurogenesis in the hippocampus. In addition, we show that two-week treatment with lithium chloride rescues long-term context fear conditioning and partially rescues hippocampal neurogenesis in *Bbs1*^M390R/M390R^ mice. Overall, this study shows a molecular connection between primary cilia and learning and memory using mouse models of BBS. Our study also identifies lithium as a potential therapeutic agent for treating the intellectual disability aspect of BBS.

## Results

### 1. *Bbs1*^*M390R/M390R*^ mice have impaired long-term fear conditioning

To study role of BBS in learning and memory, we employed a mouse model of the most common human BBS mutation, *Bbs1*^*M390R/M390R*^. Learning was evaluated using a three-day delay fear conditioning paradigm, which tests for long term association memory (Fig 1A). Controls were littermate heterozygote or wild-type mice as there is no difference in fear conditioning between these animals (S1 Fig). On day 1, both *Bbs1*^*M390R/M390R*^ mice and their littermate controls showed increased freezing behavior following a shock stimulus, indicating that BBS mice exhibit a normal physiological response to an aversive stimulus (Fig 1B). On day 2 (post 24 hours from training), mice were introduced into a novel environment to test cue (sound) dependent fear conditioning. We found no significant differences between *Bbs1*^*M390R/M390R*^ and control mice, indicating that BBS mice have intact cue dependent learning (Fig 1C). On day 3 (post 48 hours from training), mice were re-introduced back into the training environment to test context (environment) dependent learning. Remarkably, *Bbs1*^*M390R/M390R*^ mice showed a 28% reduction in freeze behavior in this environment relative to littermate controls (Fig 1C). A sex difference was not observed in control mice or *Bbs1*^*M390R/M390R*^ mice for acquisition (immediate fear conditioning), cue fear conditioning (24 hours after acquisition), and context fear conditioning (48 hours after acquisition) (S2 Fig). These findings revealed that BBS mice have context specific fear conditioning impairments.

**Fig 1.**
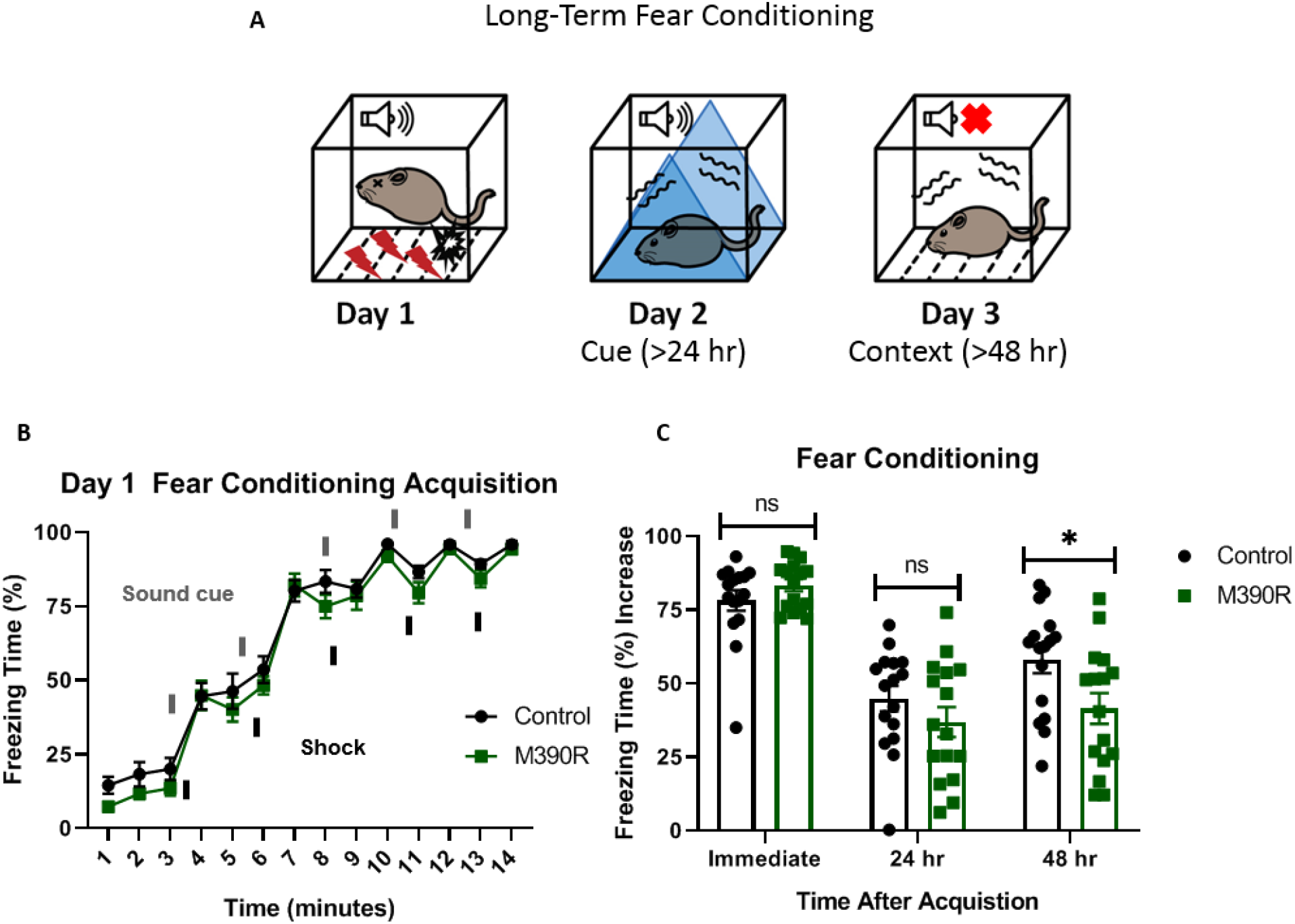
*Bbs1*^*M390R/M390R*^ mice have impaired long-term context fear conditioning. A.) Schematic diagram of the delay fear conditioning procedure. On the first day, a mouse was placed in a chamber, and a sound was paired with a shock multiple times. On the second day, a mouse was placed in an altered chamber. The chamber was triangle shaped (represented by the blue triangle) with a smooth floor, and a sound was given to test long-term cue fear conditioning. On the third day, the mouse was placed back in the same chamber (context) as day 1, without a sound cue. This set up was used to test long-term context fear conditioning. B.) Day 1 acquisition between the control mice (n=16) and the *Bbs1*^*M390R/M390R*^ mice (n=16) for long-term fear conditioning differed significantly (2-way ANOVA, time X genotype, F (13, 420) = 0.4488, P=0.950, time, F (13, 420) = 179.8, P<0.0001, genotype, F (1, 420) = 11.46, P=0.0008). The thick lines above the curve indicate when the sound cue was given, and the thick lines below the curve indicate when the shock was given. C.) The immediate fear conditioning indicates training to the day 1 fear conditioning. The immediate fear conditioning was measured as the difference of the freezing time (%) just before conditioning (first three minutes) and just after conditioning (last minute). The immediate fear conditioning did not differ significantly between the control mice (n=16) and *Bbs1*^*M390/M390R*^ mice (n=16) used for long-term fear conditioning (Welch’s t-test, P=0.2107). The post 24 hr fear conditioning represents cue fear conditioning, and is portrayed as Day 2 on the schematic diagram. The 24 hr fear conditioning (cue) was measured as the difference of the freezing time (%) before the tone (cue) on day 2 and during the tone (cue) on day 2. The 24 hr fear conditioning (cue) did not differ significantly between the control mice (n=16) and the *Bbs1*^*M390R/M390R*^ mice (n=16) (Welch’s t-test, P=0.2414). The post 48 hr fear conditioning represents context fear conditioning, and is portrayed as Day 3 on the schematic diagram. The 48 hr fear conditioning was measured as the difference of the freezing time (%) just before conditioning (first three minutes of day 1) and during the context on day 3. The 48 hr fear conditioning (context) between the control mice (n=16) and the *Bbs1*^*M390R/M390R*^ mice (n=16) differed significantly (Welch’s t-test, P=0.0240). control mice= *Bbs1*^*M390R/+*^ mice, M390R=*Bbs1*^*M390R/M390R*^ mice, hr = hour, ns = not significant * P< 0.05, ** P< 0.01, ***P<0.001 ****P<0.0001

Due to the pleiotropic nature of BBS, we tested for confounding factors that may underlie the striking impairments in fear conditioning observed in *Bbs1*^*M390R/M390R*^ mice. No hearing differences were observed between *Bbs1*^M390R/M390R^ mice and control mice based on Auditory Brainstem Response and hearing behavior (S3A and S3B Fig). No differences were observed in shock reactivity between *Bbs1*^*M390R/M390R*^ mice and control mice indicating a normal tactile response (S3C Fig). Moreover, we did not observe a difference in activity levels or sleep behavior between *Bbs1*^*M390R/M390R*^ mice and control mice (S3D and S3E Fig). These findings revealed that the impaired fear response is not due to a secondary effect of these sensory systems.

### 2. *Bbs1*^*M390R/M390R*^ mice have normal short-term fear conditioning

We tested short-term fear conditioning in BBS mice to assess if the long-term memory deficit is due to short-term memory impairment. To test for short-term fear conditioning, we used a 1-day fear conditioning paradigm (Fig 2A). Both the control mice (*Bbs1*^*M390R/+*^ mice) and *Bbs1*^*M390R/M390R*^ mice showed intact conditioning to shock (Fig 2B). In addition, there was no significant difference in short-term context memory between the control mice and *Bbs1*^*M390R/M390R*^ mice (Fig 2C). These results are in contrast to the differences in long-term context memory, which showed impaired performance in *Bbs1*^*M390R/M390R*^ mice compared to controls (Fig 1C). These findings indicated that BBS mice have specific impairments in long-term context fear conditioning.

**Fig 2.**
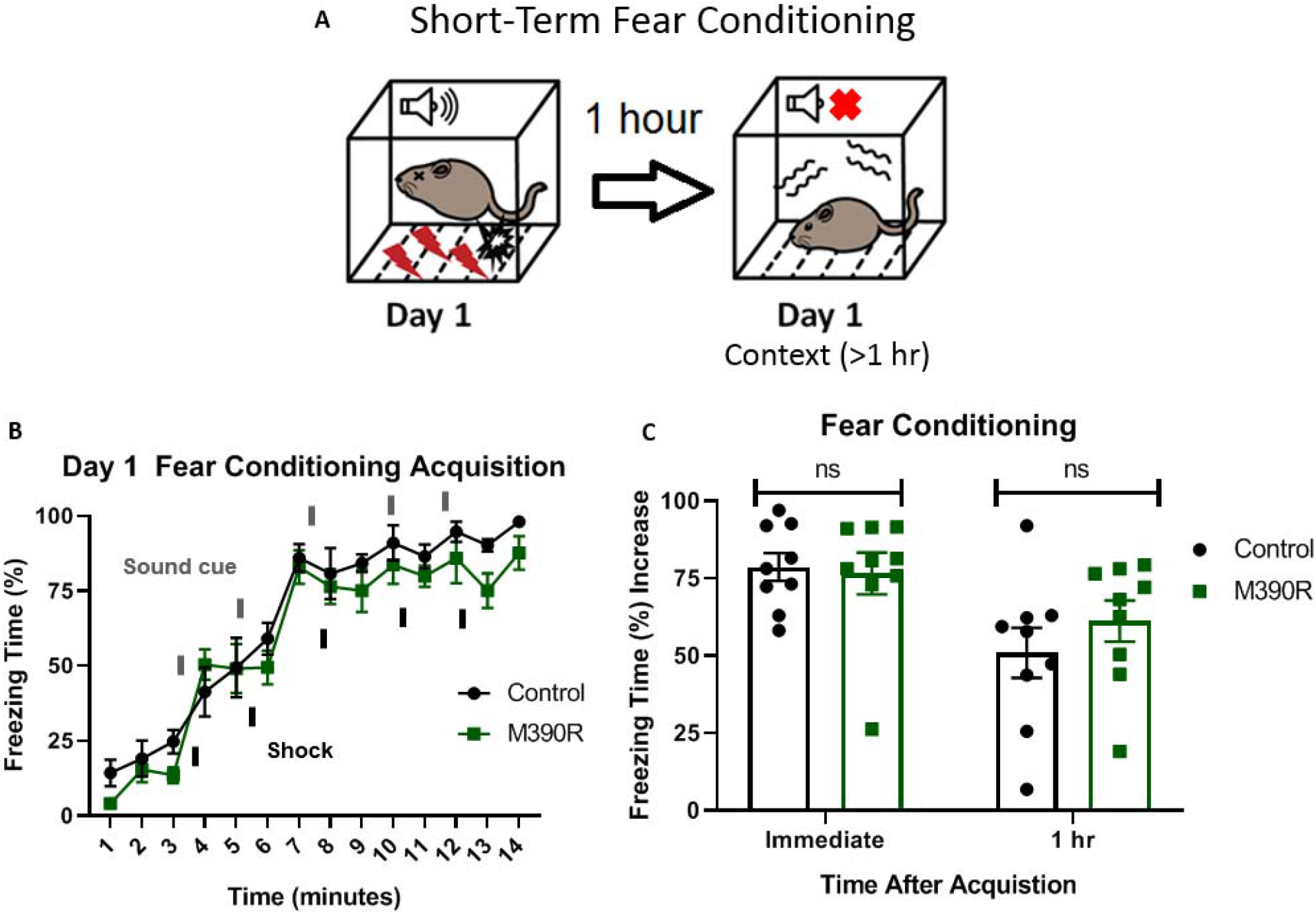
*Bbs1*^*M390R/M390R*^ mice have normal short-term context fear conditioning. A.) Schematic diagram of the one day delay fear conditioning procedure. On the first day, a mouse was placed in a chamber, and a sound was paired with a shock multiple time. One hour later, the mouse was placed back in the chamber, and freezing was measured for short-term context fear conditioning. B.) Day 1 acquisition between the control mice (n=9) and the *Bbs1*^*M390R/M390R*^ mice (n=9) used for the short term fear conditioning differed significantly (2-way ANOVA, time X genotype, F (13, 224) = 0.5574, P=0.8858, time, F (13, 224) = 56.19, P<0.0001, genotype, F (1, 224) = 9.369, P=0.0025). The thick lines above the curve indicate when the sound cue was given, and the thick lines below the curve indicate when the shock was given. C.) The immediate fear conditioning indicates training to the day 1 fear conditioning. The immediate fear conditioning was measured as the difference of the freezing time (%) just before conditioning (first three minutes) and just after conditioning (last minute). The immediate fear conditioning did not differ significantly between control mice (n=9) and *Bbs1*^*M390/M390R*^ mice (n=9) used for the short-term fear conditioning (Welch’s t-test, P=0.8004). The 1 hr fear conditioning represents short-term context fear conditioning. The 1 hr fear conditioning was measured as the difference of the freezing time (%) just before conditioning and 1 hour after conditioning. The day 1 fear conditioning for context between the control mice (n=9) and the *Bbs1*^*M390R/M390R*^ mice (n=9) did not reveal a significant difference (Welch’s t-test, P=0.3436). control mice= *Bbs1*^*M390R/+*^ mice, M390R=*Bbs1*^*M390R/M390R*^ mice, hr = hour, ns = not significant * P< 0.05, ** P< 0.01, ***P<0.001, ****P<0.0001

### 3. Mice with postnatal deletion of Bbs8 have impaired long-term context fear conditioning

We took advantage of another BBS mouse model to further explore the role of BBS genes (especially BBSome genes) in fear conditioning, specifically a tamoxifen inducible knock-out mouse model of BBS8 [36]. BBS8, like BBS1, is a component of the BBSome [11]. Using *Bbs8* tamoxifen inducible knock-out mice, we evaluated the temporal effects of BBS8 on fear conditioning (Fig 3A). For controls, we used littermates lacking *Cre.* Tamoxifen was administered to both groups of mice to control for possible effects of tamoxifen on behavior [37]. Following day 1 of fear conditioning, mice with *Bbs8* postnatally deleted, as well as control mice, were successfully conditioned to fear (Fig 3B). However, significant impairments in context but not cue fear conditioning were observed between conditional KO *Bbs8* mice and controls (Fig 3C). These results are similar to the results for *Bbs1*^*M390R/M390R*^ mice, indicating a role of the BBSome in mediating long-term context fear conditioning. Although, we observed differences in the acquisition curve of day 1 fear conditioning between conditional KO *Bbs8* mice and controls, no difference was found in immediate fear conditioning (Fig 3B and 3C). These results indicate that like *Bbs1*, *Bbs8* is involved in long-term context fear conditioning.

**Fig 3.**
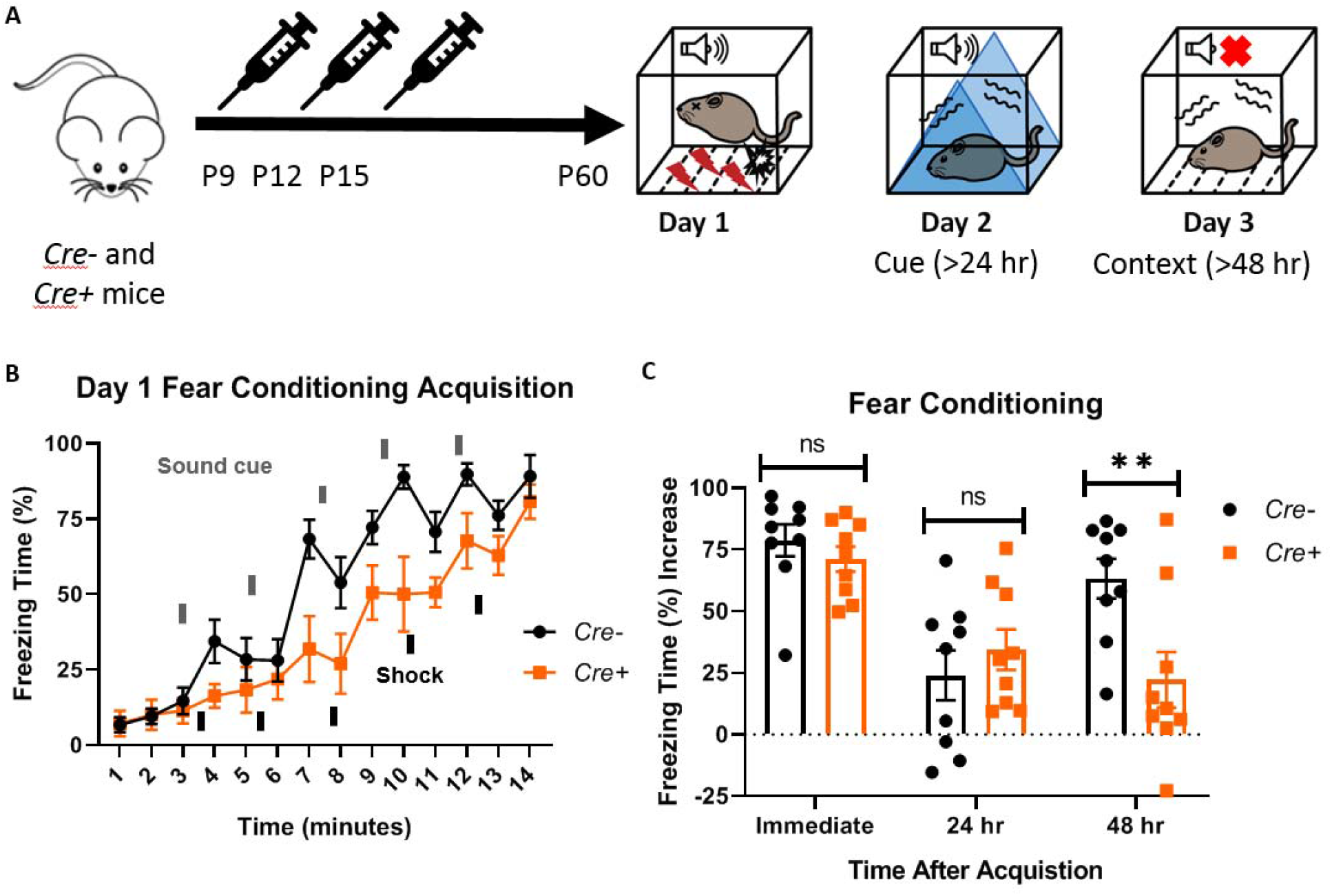
Postnatal BBS genes are involved in long-term context fear conditioning. A.) Timeline of the tamoxifen I.P injections of the experimental mice, *Cre+* (*Bbs8*^flox/−^ and *Bbs8*^flox/flox^; *UBC-Cre*^*ERT2*^) and littermate control mice, *Cre-* (*Bbs8*^flox/−^ and *Bbs8*^flox/flox^; *UBC-Cre*^*ERT2*^-). To induce *Bbs8* deletion in *Cre+* mice, Tamoxifen was injected at P9, P12, and P15 (denoted by the syringe image). At 2 months of age, the mice were tested for long-term fear conditioning. The first day was the acquisition phase for fear conditioning, the second day was cue fear conditioning, and the third day was context fear conditioning. B.) Day 1 acquisition curve between the *Cre-* mice (n=9) and *Cre+* mice (n=9) differed significantly (2-way ANOVA, time X genotype, F (13, 224) = 1.721, p=0.0579, time, F (13, 224) = 31.71, P<0.0001, genotype, F (1, 224) = 39.16, P<0.0001). The thick lines above the curve indicate when the sound cue was given, and the thick lines below the curve indicate when the shock was given. C.) The immediate fear conditioning indicates training to the day 1 fear conditioning. The immediate fear conditioning was measured as the difference of the freezing time (%) just before conditioning (first three minutes) and just after conditioning (last minute). The immediate fear conditioning did not differ significantly between the *Cre-* mice (n=9) and *Cre+* mice (n=9) used for long-term fear conditioning (Welch’s t-test, P=0.3717). The post 24 hr fear conditioning represents cue fear conditioning, and is portrayed as Day 2 on the schematic diagram. The 24 hr fear conditioning (cue) was measured as the difference of the freezing time (%) before the tone (cue) on day 2 and during the tone (cue) on day 2. The 24 hr fear conditioning (cue) did not differ significantly between the *Cre-* mice (n=9) and *Cre+* mice (n=9) (Welch’s t-test, P=0.4325). The post 48 hr fear conditioning represents context fear conditioning, and is portrayed as Day 3 on the schematic diagram. The 48 hr fear conditioning was measured as the difference of the freezing time (%) just before conditioning (first three minutes of day 1) and during the context on day 3. The 48 hr fear conditioning (context) between the *Cre-* mice (n=9) and *Cre+* mice (n=9) differed significantly (Welch’s t-test, P=0.0099). *Cre- = Bbs8*^flox/−^ and *Bbs8*^flox/flox^; *UBC-Cre*^*ERT2*^- mice, *Cre+* = *Bbs8*^flox/−^ and *Bbs8*^flox/flox^; *UBC-Cre*^*ERT2*^ + mice, hr = hour, del=deletion, flx= flox, hr=hour, ns =not significant * P< 0.05, ** P< 0.01, ***P<0.001, ****P<0.0001

### 4. Mice with preferential deletion of *Bbs1* in the forebrain have impaired long-term context fear conditioning

The forebrain contains brain regions involved in fear conditioning including the amygdala and hippocampus[30]. To explore whether the absence of normal BBS1 function in the forebrain is responsible for the fear conditioning impairment observed in *Bbs1*^*M390R/M390R*^ mice, we utilized a forebrain-specific *Bbs1* knock-out mouse line developed by crossing a *Bbs1*^*flox/flox*^ conditional mouse line with a *Cre* line expressed in the forebrain (*Emx1*-*Cre* mice). The *Emx1*-*Cre* mice were generated and verified by Gorski et al. [38]. Using an Ai9 Cre reporter allele, we confirmed that the Cre is preferentially expressed in the forebrain (S4A and S4B Fig). We also confirmed that *Bbs1* is excised in the forebrain, but unexcised in the hindbrain of *Bbs1*^*flox/flox*^, *Emx1-Cre*+ mice (S4C Fig), further confirming the specificity of *Cre* expression in *Emx1*-*Cre* mice.

Control mice (*Emx1*-*Cre, Bbs1*^*+/+*^ mice) and forebrain specific *Bbs1* knockout mice (*Emx1*-*Cre*; *Bbs1*^*flox/−*^ mice) were fear conditioned using a three-day fear conditioning paradigm (Fig 4A). Acquisition of conditioning to shock was intact for both control mice and forebrain specific *Bbs1* knockout mice (Fig 4B). However, forebrain specific *Bbs1* knockout mice showed impaired context fear conditioning compared to controls (Fig 4C). Cue fear conditioning was observed to be intact for both knockout and control mice. These results indicate that BBS1 in the forebrain is required for contextual memory.

**Fig 4.**
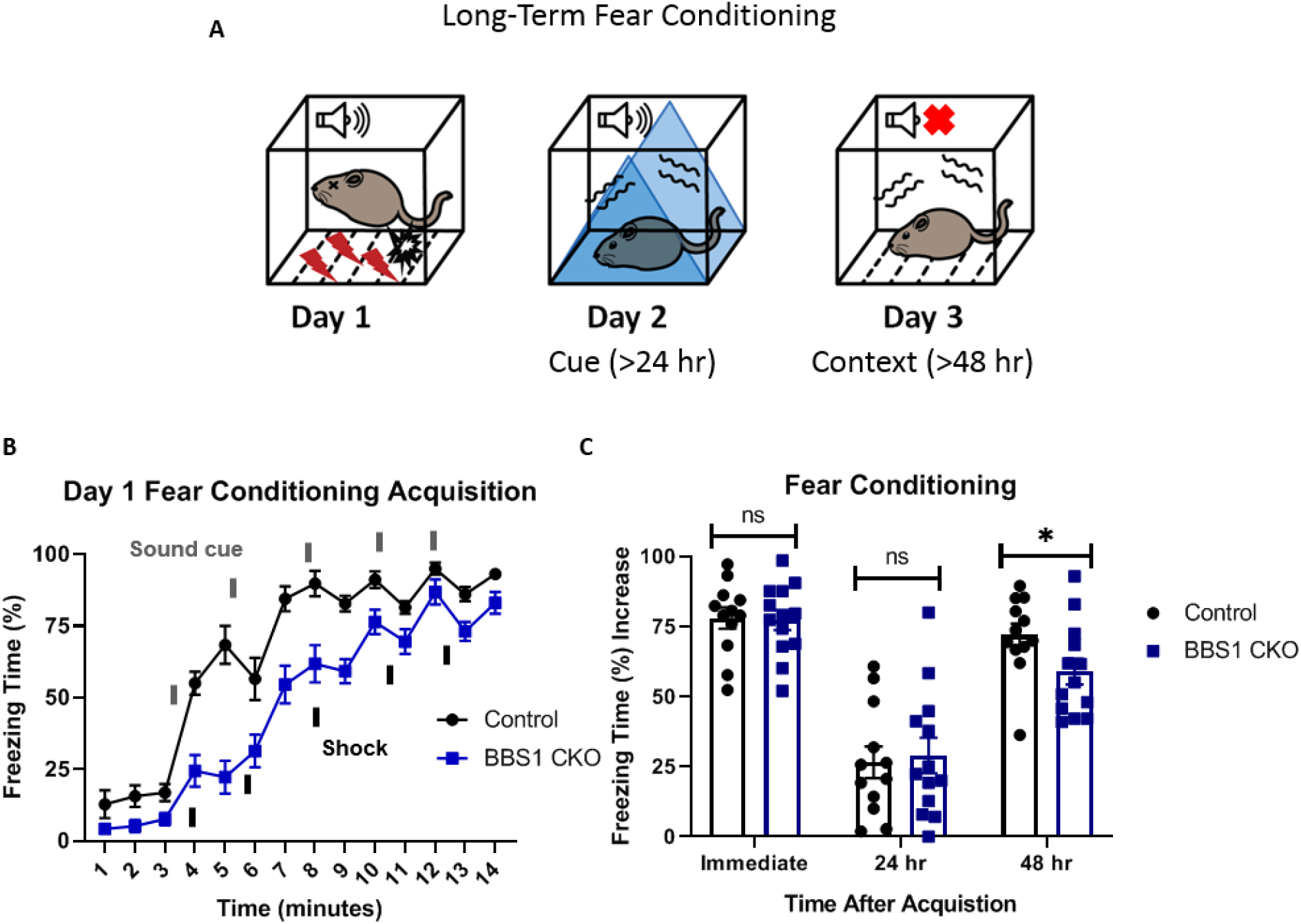
BBS genes in the forebrain are involved in long-term context fear conditioning. A.) Schematic diagram of the three day delay fear conditioning procedure. On the first day, a mouse was placed in a chamber, and a sound was paired with a shock multiple time. On the second day, a mouse was placed in an altered chamber that was triangle shaped (represented by the blue triangle) with a smooth floor, and a sound was given to measure cue fear conditioning. On the third day, the mouse was placed back in the same chamber, and measured for freezing without sound. This gives the context fear conditioning. B.) Day 1 acquisition between the *Emx1-Cre* mice (control, mixed strain of C57BL/6 and 129/SVeV, n=12) and the *Bbs1*^*flox/−*^; *Emx1-Cre* (BBS1 CKO, mixed strain of C57BL/6 and 129/SVeV, n=13) differed significantly (2-way ANOVA, time X genotype, F (13, 322) = 3.483, P<0.0001, time, F (13, 322) = 93.95, P<0.0001, genotype, F (1, 322) = 137.5, P<0.0001). The thick lines above the curve indicate when the sound cue was given, and the thick lines below the curve indicate when the shock was given. C.) The immediate fear conditioning indicates training to the day 1 fear conditioning. The immediate fear conditioning was measured as the difference of the freezing time (%) just before conditioning (first three minutes) and just after conditioning (last minute). The immediate fear conditioning did not differ significantly between the control *Emx1-Cre* mice (n=12) and *Bbs1*^*flox/−*^; *Emx1-Cre* (n=13) used for the long-term fear conditioning (Welch’s t-test, P=0.8999). The post 24 hr fear conditioning represents cue fear conditioning, and is portrayed as Day 2 on the schematic diagram. The 24 hr fear conditioning (cue) was measured as the difference of the freezing time (%) before the tone (cue) on day 2 and during the tone (cue) on day 2. The 24 hr fear conditioning (cue) did not differ significantly between the control *Emx1-Cre* mice (n=12) and *Bbs1*^*flox/−*^; *Emx1-Cre* (n=13) (Welch’s t-test, P=0.7005). The post 48 hr fear conditioning represents context fear conditioning, and is portrayed as Day 3 on the schematic diagram. The 48 hr fear conditioning was measured as the difference of the freezing time (%) just before conditioning (first three minutes of day 1) and during the context on day 3. Day 3 fear conditioning for context between the control *Emx1-Cre* mice (n=12) and *Bbs1*^*flox/−*^; *Emx1-Cre* (n=13) differed significantly (Welch’s t-test, P=0.0438, Mann-Whitney-Wilcoxon Test, P=0.0398). control = *Bbs1*^*+/+*^; *Emx1*-Cre mice, BBS1 CKO = *Bbs1*^*flox/−*^; *Emx1-Cre+* mice, hr=hour, ns =not significant * P< 0.05, ** P< 0.01, ***P<0.001, ****P<0.0001

### 5. *Bbs1*^M390R/M390R^ mice do not have impaired long-term potentiation

Since *Bbs1*^*M390R/M390R*^ mice have impaired context fear conditioning, which is hippocampus dependent [30], we investigated hippocampal function in *Bbs1*^*M390R/M390R*^ mice. We evaluated long-term potentiation (LTP) in the CA1 region of the hippocampus because LTP is a neural correlate for long-term memory consolidation [39]. In addition, some mouse models with impaired context fear conditioning have impaired long-term potentiation (LTP) in CA1 of the hippocampus [32, 40, 41]. Despite the important role of LTP in fear conditioning, we did not observe a difference in LTP between control mice and *Bbs1*^*M390R/M390R*^ mice in CA1 of the hippocampus (Fig 5A and 5B). These results suggest that the observed impaired learning arises from causes other than impaired LTP.

**Fig 5.**
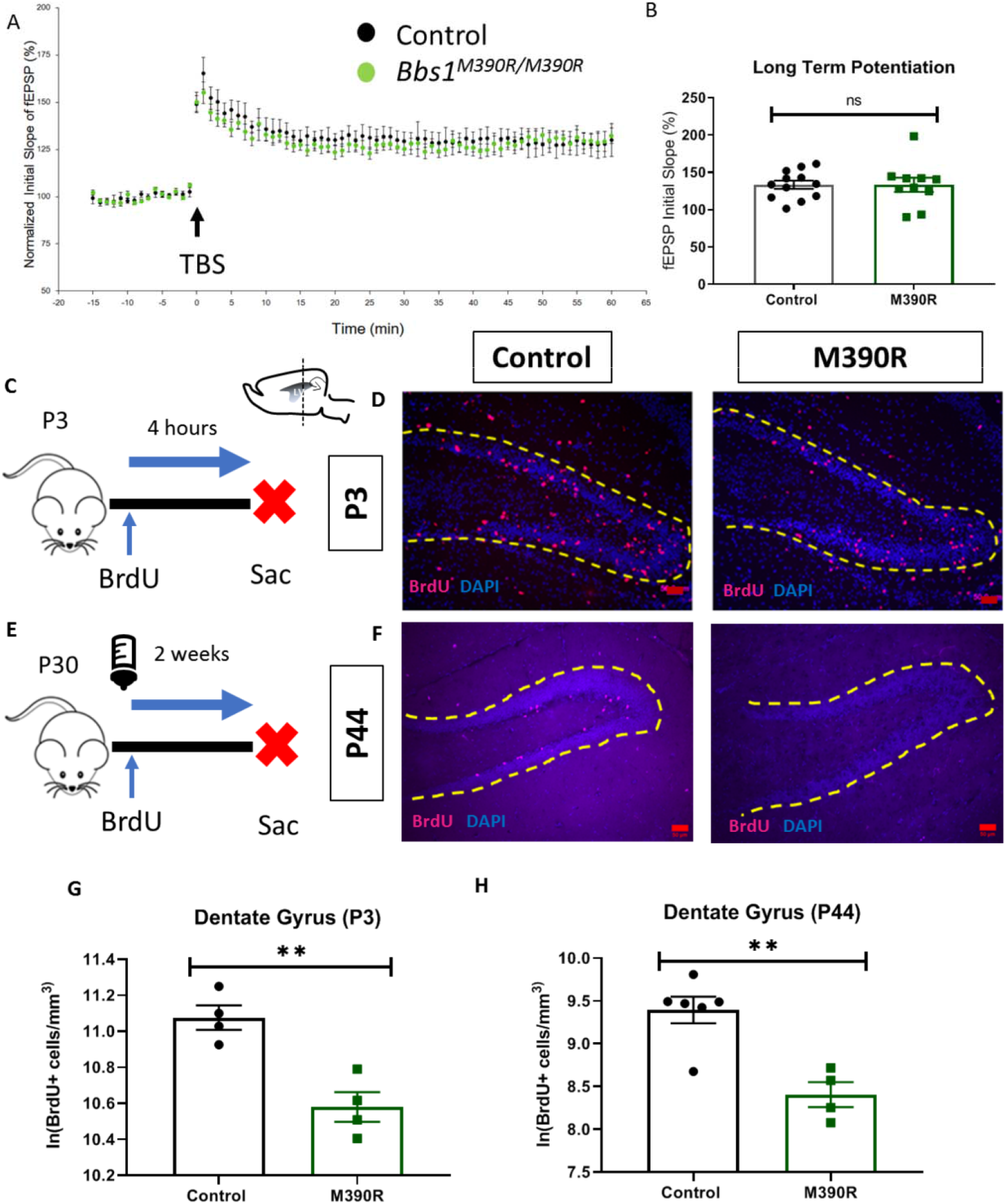
*Bbs1*^*M390R/M390R*^ mice have decreased hippocampal proliferation. A.) Normalized initial slope (%) recordings of field excitatory post synaptic potentials (fEPSP) in the hippocampal CA1 Schaffer-collateral pathway between 2 month male control mice (n=17, 4 mice) and *Bbs1*^*M390R/M390R*^ mice (n=16, 4 mice). LTP was induced by 12 theta burst stimulation (TBS). B.) The Long Term Potentiation (average of last five minutes of normalized initial slope of fEPSP) in the hippocampal CA1 Schaffer-collateral pathway between 2 month male control mice (n=17, 4 mice) and *Bbs1*^*M390R/M390R*^ mice (n=16, 4 mice) did not differ significantly (Welch’s t-test, P=0.8407). C.) Schematic diagram of the BrdU injections of postnatal day 3 (P3) mice. P3 mice were IP injected with 300mg/kg BrdU, and taken down four hours later. Sac=Sacrifice D.) Inverted fluorescent microscope images of the P3 Dentate Gyrus. The sections were stained with Bromodeoxyruidine (BrdU) and counterstained with the nuclear marker, DAPI. The yellow dotted line outlines the dentate gyrus. The Red Bar line was 50um. E.) Schematic diagram of the BrdU procedures for postnatal day 44 (P44) mice. At P30, mice were started on BrdU injections (2×50mg/kg) for five days. At P44, mice were taken down. Sac=Sacrifice F.) Inverted fluorescent microscope images of the P44 Dentate Gyrus. The sections were stained with Bromodeoxyruidine (BrdU) and counterstained with the nuclear marker, DAPI. The yellow dotted line outlines the dentate gyrus. The Red Bar line was 50um. G.) Decreased proliferation in the dentate gyrus of P3 *Bbs1*^*M390R/M390R*^ mice. The natural logarithm of BrdU+cell/mm^3^ in the dentate gyrus between the control mice (n=4) and the *Bbs1*^*M390R/M390R*^ mice (n=4) differed significantly (Welch’s t-test, P=0.0038). H.) Decreased proliferation in the dentate gyrus of P44 *Bbs1*^*M390R/M390R*^ mice. The natural logarithm of BrdU+cell/mm^3^ in the dentate gyrus between the control mice (n=6) and the *Bbs1*^*M390R/M390R*^ mice (n=4) differed significantly (Welch’s t-test, P=0.0018). control=*Bbs1*^*+/+*^, *Bbs1*^*M390R/+*^ mice, M390R=*Bbs1*^*M390R/M390R*^ mice, hr=hour, ns=not significant, * P< 0.05, ** P< 0.01, ***P<0.001, ****P<0.0001

### 6. *Bbs1*^M390R/M390R^ mice have decreased hippocampal neurogenesis

Next, we sought to identify a potential cause of the defective long-term fear conditioning observed in BBS mice, It has been recently reported that BBS patients have decreased hippocampal volume which is thought to be a result of impaired neurogenesis [42]. Due to the known role that cilia play in mediating cell proliferation and hippocampal volume in patients [43, 44], we hypothesized that defective hippocampal neurogenesis underlies the fear conditioning deficits in BBS mice. Therefore, we investigated hippocampal neurogenesis in *Bbs1*^*M390R/M390R*^ mice.

To measure proliferation, we injected *Bbs1*^*M390R/M390R*^ mice and control mice with BrdU, a thymidine analog that is incorporated into replicating DNA to label proliferating cells (Fig 5C and 5E). Following BrdU labeling, we found that both neonatal (P3) and young adult (P44) *Bbs1*^*M390R/M390R*^ mice displayed significant reductions in BrdU+ cells in the hippocampal dentate gyrus compared to controls (Fig 5D, Fig 5F-5H). Moreover, young adult *Bbs1*^M390R/M390R^ mice also showed fewer new neurons as determined by a reduced number of cells co-labeled for BrdU and Doublecortin, a marker for immature neurons(Fig 6B and 6D) [45].

**Fig 6.**
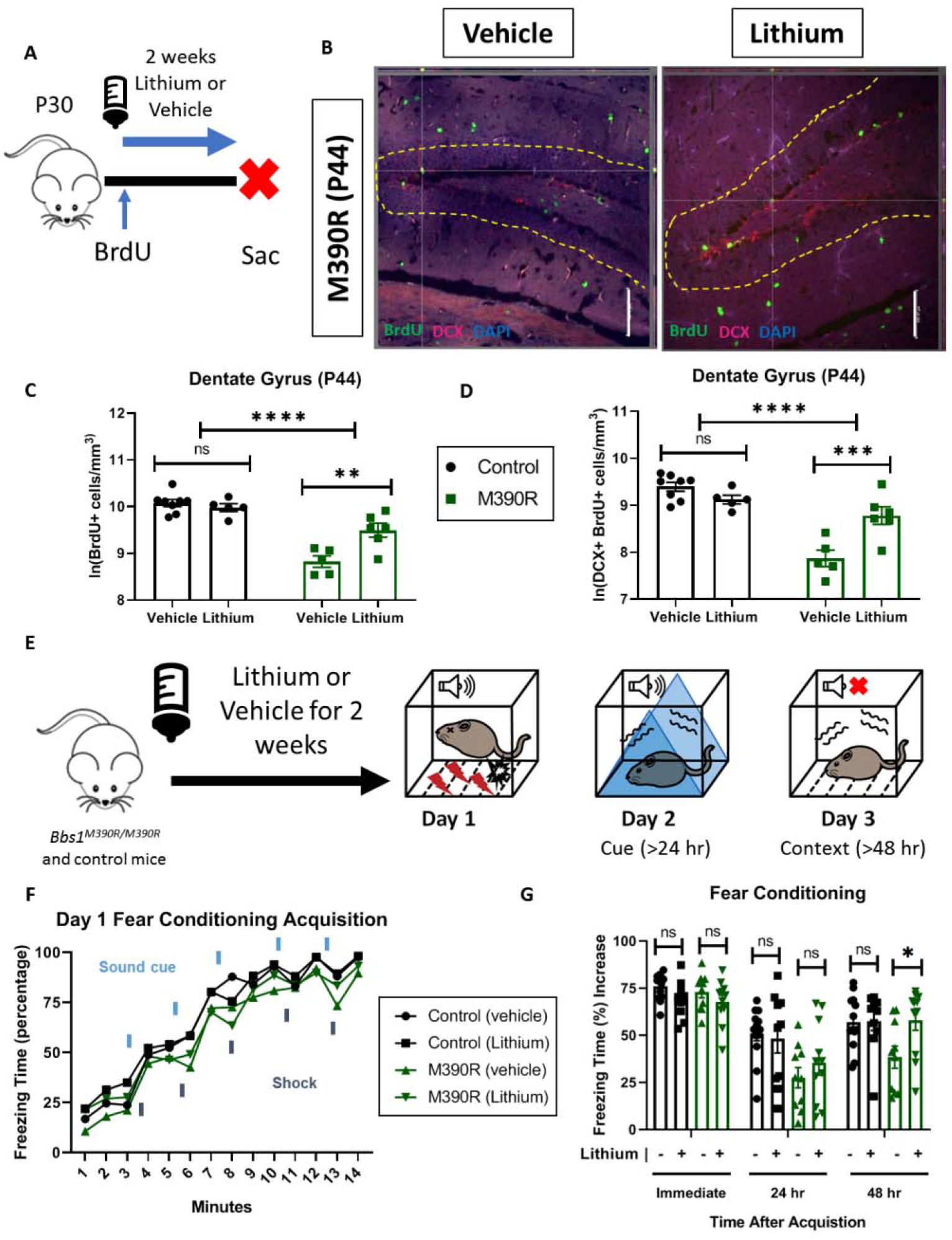
Chronic Lithium treatment rescued long-term context fear conditioning in *Bbs1*^*M390R/M390R*^ mice. A.) Schematic diagram of the Bromodeoxyuridine (BrdU) procedures for postnatal day 44 (P44) mice. At P30, mice were started on Lithium water (45mM) or continued with water (vehicle). Mice were also started on BrdU injections (2×50mg/kg) for five days. At P44, mice were taken down. Sac=Sacrifice B.) Images of immunohistochemistry for neurogenesis of vehicle and lithium treated *Bbs1*^*M390R/M390R*^ mice. Z-stack, 20X, images of Dentate Gyrus at postnatal day 44 (P44). The tissue sections were stained with BrdU, Doublecortin (DCX), and counterstained with the nuclear marker DAPI. The yellow dotted line outlines the dentate gyrus. The White Bar line indicates 100micrometers. C.) Proliferation in the dentate gyrus of P44 control and *Bbs1*^*M390R/M390R*^ mice. We assessed two factors, and found a significant interaction for genotype and treatment, and a difference in treatment and genotype (2-way ANOVA, treatment X genotype, F (1, 20) = 6.428, P=0.0026, treatment, F (1, 20) = 1.569, P=0.0177, genotype, F (1, 20) = 46.89, P<0.0001). A Sidak’s multiple comparisons test showed a significant difference in treatment for *Bbs1*^*M390R/M390R*^ mice (n=5 Vehicle, n=6 Lithium, P=0.7875) but not for control mice (n=8 Vehicle, n=5 Lithium, P=0.0010) D.) Neurogenesis in the dentate gyrus of P44 control and *Bbs1*^*M390R/M390R*^ mice. We assessed two factors, and found a significant interaction between genotype and treatment, and a difference in treatment and genotype (2-way ANOVA, treatment X genotype, F (1, 20) = 11.37, P=0.0005, treatment, F (1, 20) = 0.0368, P=0.5779, genotype, F (1, 20) = 29.54, P<0.0001). A Sidak’s multiple comparisons test showed a significant difference in treatment for *Bbs1*^*M390R/M390R*^ mice (n=5 Vehicle, n=6 Lithium, P=0.0006) but not for control mice (n=8 Vehicle, n=5 Lithium, P=0.3301). E.) Timeline of LiCl treatment. At 4-5 weeks of age, mice were treated with LiCl (45mmol) water or continued on water (vehicle). After two weeks of treatment, mice were tested on a 3 day fear conditioning set up. Day 1 was the training (acquisition phase) for fear conditioning. The second day was testing for cue fear conditioning. The third day was testing for context fear conditioning. F.) Day 1 fear conditioning acquisition between the vehicle treated control mice (n=13) and *Bbs1*^*M390R/M390R*^ (45mmol) treated mice (n=10), and lithium treated control mice (n=9) and *Bbs1*^*M390R/M390R*^ (45mmol) treated mice (n=11). Graph was presented without standard error. We found a significant difference in Treatment, Genotype, and Time (3-way ANOVA, Time x Genotype x Treatment, F (13, 560) = 0.2572, P=0.9963; Genotype x Treatment, F (1, 560) = 0.4214, P=0.5165; Time x Treatment, F (13, 560) = 1.338, P=0.1860; Time x Genotype, F (13, 560) = 0.5960, P=0.8582; Treatment, F (1, 560) = 5.475, P=0.0196; Genotype, F (1, 560) = 39.48, P<0.0001; Time, F (13, 560) = 157.3, P<0.0001). The thick lines above the curve indicate when the sound cue was given, and the thick lines below the curve indicate when the shock was given. G.) The immediate fear conditioning indicates training to the day 1 fear conditioning. The immediate fear conditioning was measured as the freezing time (%) increase of the freezing time (%) just after conditioning (last minute) to the freezing time (%) just before conditioning (first three minutes). The immediate fear conditioning was not significantly different between the control mice given vehicle (n=13) and control mice given LiCl water (n=10) (Welch’s t-test, P=0.0689) and did not significantly differ between the *Bbs1*^*M390R/M390R*^ mice given vehicle (n=10) and *Bbs1*^*M390R/M390R*^ mice given LiCl water (45mmol) (n=11) (Welch’s t-test, P=0.2834). The post 24 hr fear conditioning represents the cue fear conditioning. The 24 hr fear conditioning (cue) was measured as the freezing time (%) increase of the freezing time (%) during the tone (cue, day 2) to the freezing time (%) before the tone (cue, day 2). The 24 hr fear conditioning (cue) was not significantly different between the control mice given vehicle (n=13) and control mice given LiCl water (n=10) (Welch’s t-test, P=0.7483) and was not significantly different between the *Bbs1*^*M390R/M390R*^ mice given vehicle (n=10) and *Bbs1*^*M390R/M390R*^ mice given LiCl water (45mmol) (n=11) (Welch’s t-test, P=0.3625). The post 48 hr fear conditioning represents the context fear conditioning. The 48 hr fear conditioning was measured as the freezing time (%) increase of the freezing time (%) during the context on day 3 to the freezing time (%) just before conditioning (first three minutes of day 1). Day 3 fear conditioning for context was not significantly different between the control mice given vehicle (n=13) and control mice given LiCl water (n=10) (Welch’s t-test with Bonferroni correction, P=0.9285) but was significantly different between the *Bbs1*^*M390R/M390R*^ mice given vehicle (n=10) and *Bbs1*^*M390R/M390R*^ mice given LiCl water(45mmol) (n=11) (Welch’s t-test, P=0.0235). control=*Bbs1*^*+/+*^, *Bbs1*^*M390R/+*^ mice, M390R=*Bbs1*^*M390R/M390R*^ mice, hr=hour, ns=not significant, * P< 0.05, ** P< 0.01, ***P<0.001, ****P<0.0001

The role of the observed impairments in hippocampal neurogenesis in long-term context fear conditioning of *Bbs1*^*M390R/M390R*^ is unclear. To test the role of neurogenesis, we utilized a pharmacological modality to enhance hippocampal neurogenesis. Because impaired neurogenesis within the dentate gyrus is associated with long term memory deficits, we reasoned that rescue of impaired neurogenesis could improve fear conditioning impairments. To this end, we chose lithium due to its previous use as an agent to improve neurogenesis and hippocampal dependent memory [46–48].

We began by assessing the effects of lithium on hippocampal neurogenesis in *Bbs1*^*M390R/M390R*^ mice and control mice. Young adult mice were treated with lithium or vehicle (water) for two weeks, and then brain tissues were harvested and stained for BrdU and Doublecortin (Fig 6A and 6B). Compared to vehicle treated mice, lithium treatment led to a 153% increase in the number of new neurons in the dentate gyrus of the hippocampus (Fig 6C and 6D).

### 7. Lithium treatment rescued long-term context fear conditioning in *Bbs1*^M390R/M390R^ mice

We hypothesized that enhancing hippocampal neurogenesis using lithium treatment would rescue context which is hippocampus dependent [30]. To test the effects of lithium on fear conditioning, 4-5 week old mice were administered lithium or vehicle for two weeks (Fig 6E). The mice underwent fear conditioning using the three-day paradigm (Fig 6E-6G). As hypothesized, lithium treatment rescued the long-term context fear conditioning, but not long-term cue fear conditioning for *Bbs1*^*M390R/M390R*^ mice (Fig 6G). Taken together, these obserations suggest BBS genes play an important role in mediating hippocampal neurogenesis and long-term context fear conditioning.

## Discussion

Intellectual disability (ID) is the most common neurodevelopmental disorder [1]. ID has limited pharmacological treatments, which is attributed to a limited understanding of the mechanisms involved. A large reason for the lack of mechanistic understanding is due to a lack of mouse models with ID as a primary phenotype. To overcome this hurdle, we explored the use of mouse models of the human disorder, Bardet-Biedl Syndrome (BBS) for the study of ID.

We showed that BBS mice have impaired context fear conditioning, indicating that BBS genes play a critical role in long-term memory. Our studies elucidated the spatial and temporal role of BBS gene function in fear memory. Using a conditional *Bbs1* knock-out mouse model, we demonstrated that BBS1 in the forebrain plays an important role in long-term fear memory. These findings are consistent with prior reports that cilia in the forebrain are involved in long-term fear memory[49]. In addition, the use of a novel tamoxifen inducible *Bbs8* knock-out mouse model demonstrated that BBS gene function is also critical during the post-natal consolidation of long-term fear memory.

Our work is in contrast to previous work using BBS mice to study their role in fear conditioning, which gave inconsistent results [50, 51]. This is partially explained by the use of different mouse strains and testing parameters. This previous work used *Bbs4* null mice on C57BL/6 [51] or FVB/NJ[50] backgrounds. In addition, these studies have noted a sex difference in BBS mice (*Bbs4* knock-out mice) with respect to fear conditioning, which we do not observe [51]. The discrepancies in findings may be due to differences in mouse strains or study design. Our study primarily used strains 129/SvEv and C57BL/6. We used a strong learning paradigm with five pairings of shocks, compared to three shock pairings[51] or two shock pairings[50]. BBS4 mice were evaluated in the previous study compared to BBS1M390R and BBS8 knock-out mice in the current study. Although BBS4, BBS1 and BBS8 are all components of the BBSome, it is possible that these proteins could have unique properties on fear memory.

Since BBS is a pleiotropic disorder, there are other factors that could explain the context fear conditioning impairment observed in mouse models of BBS. The *Bbs1*^*M390R/M390R*^ mice have visual deficits [24], olfactory deficits [52], obesity [24] and hydrocephalus, which could globally affect fear conditioning. In order to control for these phenotypes, we used young adult mice prior to the onset of obesity and blindness. In addition, our BBS1 conditional knock-out mice are not blind nor obese and BBS8 conditional knock-out mice do not have hydrocephalus [19, 53], yet both models have impaired long-term context fear conditioning. We are not able to account for the olfactory deficit as a confounding factor. However, if these phenotypes underlie the observed fear conditioning deficits, *Bbs1*^*M390R/M390R*^ mice would also display short-term (immediate) fear conditioning deficits in addition to long-term deficits, which we did not observe. Therefore, we conclude that the fear learning deficits observed are a primary phenotype due to the absence of BBS gene function.

Other mouse models demonstrate fear memory deficits similar to those we report in this study. For example, long-term context fear conditioning, but not short-term context fear conditioning, has been reported in mice with absent neuronal nitric oxide synthase [34], mice with inhibited protein synthesis [33], and in mice with PKA [32, 33] or MAP Kinase deficiencies [33]. While these mouse models have impaired long-term potentiation (LTP) in CA1 of the hippocampus, there are mouse models with impaired memory that have normal LTP [49, 54] as is the case with our *Bbs1*^*M390R/M390R*^ mice.

The decreased hippocampal neurogenesis in *Bbs1*^*M390R/M390R*^ mice is a novel finding that can explain the impaired fear context memory. Hippocampal neurogenesis is involved in hippocampus dependent learning, such as context fear conditioning. Impaired context fear conditioning has been reported in mice with genetic suppression of proliferation of GFAP expressing cells [55, 56] and Nestin expressing cells [57, 58]. Impaired fear context long-term memory has also been repoted in mice with suppressed hippocampal neurogenesis through irradiation of the head [59] and ganciclovir treated mice [55], supporting our results using BBS mouse models.

We speculate that BBS proteins affect hippocampal neurogenesis because BBS is involved in ciliary receptor trafficking of the Smoothened Receptor [13, 60, 61], which is involved in in SHH signaling. SHH signaling is mediated by primary cilia [43]. Primary cilia are particularly enriched in the hippocampus [62]. Furthermore, SHH signaling has a proliferative effect on adult hippocampal progenitors *in vitro and in vivo* [63]. In addition, both primary cilia and smoothened receptors (hedgehog signaling) are required by adult neural stem cells [64]. The role of BBS in hippocampal proliferation may also be due to their involvement in tyrosine receptor kinase B (TrkB) receptor signaling [17]. Brain derived neurotrophic factor (BDNF) has been shown to increase neurogenesis through TrkB receptors [65, 66].

We investigated lithium as a treatment for the memory and neural deficits of *Bbs1*^*M390R/M390R*^ mice. Lithium has been shown to improve learning and memory tasks in mouse models of cognitive disease including Fragile X syndrome[67], Down syndrome[48], and Alzheimer disease[46]. In addition, lithium treatment affects the morphology of primary cilia in the brain[68] and lithium has also been shown to increase hippocampal neurogenesis [46, 48, 69]. While lithium treatment of *Bbs1*^*M390R/M390R*^ mice produced a robust effect on memory performance, lithium treatment produced a more modest response in hippocampal proliferation and neurogenesis. This suggests that a modest change in hippocampal proliferation and neurogenesis can produce a profound effect on memory. Consistent with these findings, a Danish study shows a correlation between lower incidence of dementia and long-term exposure to lithium in drinking water[70]. In addition, a study in China showed that low-dose lithium treatment improved cognitive performance in children with ID without major side effects[71]. Our results demonstrate the therapeutic potential of an FDA approved drug, lithium, for treating the cognitive and neural phenotypes of BBS patients.

It is possible that lithium has other neural effects contributing to the rescue of context fear conditioning. For example, lithium has been reported to alter dendritic spine density in the hippocampus [72] and to improve olfaction in mouse models of olfactory impairment [73, 74].

Further research is needed to explore factors involved in decreased hippocampal neurogenesis in *Bbs1*^*M390R/M390R*^ mice. In our young adult mouse study, we observed a decrease in the number of BrdU+/Doublecortin+ cells in the hippocampus, indicating decreased neurogenesis. However, the apparent decrease in neurogenesis could be due to decreased proliferation, decreased differentiation and/or decreased. Any of these factors could be likely because BBS proteins are involved in the function of primary cilia, and primary cilia are involved in proliferation, differentiation [75] and survival[76].

There are alternative explanations for the cause of the impaired learning and memory in BBS mouse models. BBS proteins traffic other ciliary receptors that are involved in learning and memory. *Bbs4* knock-out mice [16] and *Bbs7* knock-out mice [77] accumulate dopamine 1 (D1) receptors in cilia. D1 receptors are involved in learning and memory[78, 79]. BBS proteins are also involved in trafficking of the somatostatin receptor 3 (SSTR3) [60] and melanin concentrating hormone receptor 1 (MCHR1) [15], both of which are involved in learning and memory [80, 81].

Our mouse model of a ciliopathy with ID robustly presented with impaired fear memory in context fear conditioning and decreased neurogenesis. In addition, our mouse model of BBS presented similarly to the mouse model of Fragile X Syndrome[82]. Fragile X syndrome is one of the most commonly inherited disorders for intellectual disability [83], and has recently been found to have defective cilia [84]. Overall, the findings presented here support the use of BBS mice as a model for ID and support the use of pro-neurogenic treatments as a possible treatment for ID.

## Methods

### Study Approval

This research was conducted in strict accordance to the Guide for the Care and Use of Laboratory Animals, 8^th^ edition, from the National Research Council. All mice were handled based on approved Institutional Animal Care and Use Committee (IACUC) protocols (#5061426 and #8072147) at the University of Iowa. Animals were housed in facilities, maintained by the Office of Animal Resources that adhere to IACUC recommendations. Mice were euthanized either by anesthesia induced by I.P injection of ketamine/xylazine followed by transcardiac perfusion, or carbon dioxide inhalation followed by cervical dislocation. Every effort was made to minimize suffering in the mice, and humane endpoints were stringently observed.

### Animals

All mice were group housed on a set 12 hr light-dark cycle and given standard chow (LM-485; Teklab, Madison, WI, USA) and water ad libitum. Mice were generated at the University of Iowa Carver College of Medicine and all experiments were performed in accordance with the Institute for Animal Care and Use Committee at the University of Iowa. For all testing, we used young adult mice (1.5-3 month old mice), unless otherwise noted. The ages of mice were chosen to keep the weight and visual processing differences between *Bbs1*^*M390R/M390R*^ and control mice to a minimal [24]. All testing was conducted during the light cycle, unless otherwise noted.

We used several strains of mice as listed below. Control mice were of the same genetic strains as the mice with which they were compared. We used male and female mice on a pure 129/SvEv genetic background for *Bbs1*^*M390R/M390R*^ mice and littermate controls (*Bbs1*^*+/+*^ and *Bbs1*^*M390R/+*^). Heterozygote mice (*Bbs1*^*M390R/+*^) do not exhibit BBS phenotypes [24, 47], and are not significantly different in fear conditioning compared to *Bbs1*^*+/+*^ mice. To generate mice with preferential *Bbs1* deletion in the forebrain, we crossed *Bbs1*^*flox/flox*^ mice (129/SvEv) [47] with *Emx1-Cre* knock-in mice (C57BL/6) (Jackson Laboratory, #005628). To verify forebrain *Cre* expression, we crossed *Emx1-Cre* knock-in mice with the Ai9 *Cre* reporter line *Gt(ROSA)26Sor*^*tm9(CAG-tdTomato)Hze*^ (C57BL/6) (Jackson Laboratory #007909). We also used conditional *Bbs8* knock-out mice (*Bbs8*^flox/flox^, C57BL/6) [36] crossed with tamoxifen-inducible *Cre* recombinase mice, B6.Cg-*Ndor1*^*Tg(UBC-cre/ERT2)1Ejb*^/2J (Jackson Laboratory #008085).

### Tamoxifen-inducible excision of *Bbs8*

We postnatally excised *Bbs8* according to previously described procedures [36]. To induce Cre expression, *Bbs8*^flox/flox^ and *Bbs8*^flox/−^; *UBC-Cre^ERT2^*+ mice were injected subcutaneously with 40 μL of tamoxifen (15 mg/mL in corn oil) on three separate days (P9, P12, and P15). *Bbs8*^flox/flox^ and *Bbs8*^flox/−^; *UBC-Cre*^*ERT2*^-mice injected with tamoxifen were the littermate controls. We assessed excision efficiency as previously described [36]. The *Bbs8* tamoxifen inducible knock-out mice (*Bbs8*^flox/flox^ and *Bbs8*^flox/−^; *UBC-Cre*^*ERT2*^+) that were determined to have less than 90% excision were excluded from the research study. This was decided as an exclusion criterion prior to conducting the study. No other mice were excluded from the research study.

### Behavioral Testing

All behavioral testing was conducted during the light cycle, unless otherwise noted.

#### Delay Fear Conditioning

For fear conditioning, mice were placed in a fear conditioning chamber with near-infrared video. Freezing was scored with the VideoFreeze software (Med Associates, St. Albans, VT, USA). Fear conditioning can distinguish short-term context memory from long-term context memory based on when context fear conditioning is tested (short-term is 1 hour after conditioning, and long-term is ≥24 hours after conditioning) [32–34]. A 3-day protocol was used to assess both long-term cue and contextual fear conditioning.

- On the first day of fear conditioning, a 20‐second tone (75◻dB) was played, which co-terminated with a 1‐second foot shock (0.75◻mA). The tone‐shock pairings occurred five times, with the shocks at 3:20m, 5:40m, 8m, 10:20m and 12:40m. For the acquisition curve figure, the freezing data was reported as the percent time the mouse was immobile for each one-minute bout. In addition, the training in day 1 fear conditioning was measured as:

○ Immediate fear conditioning = freezing time (%) just after conditioning (last minute) - the freezing time (%) just before conditioning (first three minutes)
- On the second day, to test cue fear conditioning, mice were tested in a novel context in which floor texture, odor, and shape of the chamber had been altered. After 3LJminutes in the chamber, a 3◻minute tone (75◻dB) was delivered, followed by an additional 4◻minutes without the tone. The cue fear conditioning was measured as:

○ Cue fear conditioning = freezing time (%) during the tone on day 2 – freezing time (%) before the tone on day 2
- On the third day, to test contextual fear conditioning, the chamber was set back to the original training context. Mice were place in the chamber for 5◻minutes. The context fear conditioning was measured as:

○ Context fear conditioning = freezing time (%) on day 3 - freezing time (%) just before conditioning (first three minutes of day 1).

#### One day fear conditioning

The acquisition protocol for three-day fear conditioning was used for the one day fear conditioning protocol. After the acquisition phase, mice were placed back into their home cage. One hour after the fear conditioning, mice were placed back into the original training chamber, and recorded for five minutes. The short-term context fear conditioning was measured as the difference of the freezing time (%) just before conditioning (first three minutes of day 1) and during the context on day 1 (one hour after fear conditioning).

#### Preyer Reflex

The Preyer reflex is the startle response to auditory stimuli. Mice were given an auditory stimulus (hand clap) in their home cage. A positive sign was noted if the mouse had a rapid movement of the whole body after the auditory stimulus.

#### Circling Behavior

Circling behavior is noted in animal models of deafness [85]. Mice were observed for 5 minutes in their home cage for circling behavior. A positive circling behavior was noted if the mouse tightly circled around itself more than two times.

### Auditory Brainstem Response

The auditory brainstem response (ABR) test provides information about the auditory sensitivity of the subject. The ABR test was conducted on 2-month old control mice (n=3) and *Bbs1*^*M390R/M390R*^ mice (n=4). The experimenter was masked to the genotype. The ABR test were conducted as previously described [86]. Briefly, clicks and tone-bursts were delivered to the testing ear through a plastic acoustic tube in a sound attenuated room. ABRs were measured using an Etymotic Research ER10B+ probe microphone (Etymotic Research, Elk Grove, IL, USA) coupled to two Tucker-Davis Technologies MF1 multi-field magnetic speakers (Tucker-Davis Technologies, Alachua, FL, USA). Click and tone-burst stimuli were presented and recorded using custom software running on a PC connected to a 24-bit external sound card (Motu UltraLite mk3, Cambridge MA, USA). A custom-built differential amplifier with a gain of 1,000 dB amplified acoustic ABR responses. Output was passed through 6-pole Butterworth high-pass (100 Hz) and low-pass (3 kHz) filters and then to a 16-bit analog-to-digital converter (100,000 sample/s). The tone bursts were 3 ms in length, in addition to 1 ms onset and offset ramps (raised cosine shape) centered at 4, 8, 16, 24, and 32 kHz. Responses were recorded using standard signal-averaging techniques for 500 or 1000 sweeps. Hearing thresholds (db SPL) were determined by decreasing the sound intensity by 5 and/or 10 db decrements and recording the lowest sound intensity level resulting in a recognizable and reproducible ABR response wave pattern. Maximum ABR thresholds were capped at 100 db SPL.

### BrdU injections

For early postnatal time points, mice were injected intraperitoneally with 300mg/kg Bromodeoxyuridine (BrdU, Sigma, St. Louis, Missouri) and sacrificed 4 hours after the injection. For later postnatal time points, mice were injected intraperitoneally with 50mg/kg BrdU twice a day for five days, and sacrificed ten days later.

### Lithium Treatment

We treated mice with 45mmol lithium chloride (Sigma) in drinking water starting at four to five weeks of age for 2 weeks. Mice were group-housed for lithium treatment and group-housed for vehicle (water) treatment. Littermate controls were group-housed.

### Tissue collections and histology

For early postnatal tissue, fresh brain tissues were collected and embedded in Optimal Cutting Temperature compound (OCT, Tissue Tek, Sakura Finetek). Eight μm sections were cut on a cryostat. For late postnatal time points, mice were anesthetized by intraperitoneal injection of Ketamine (17.5 mg/cc)/Xylazine (2.5 mg/cc) at 100 μL/20 gram body weight, and transcardially perfused with 4% paraformaldehyde in phosphate-buffered saline (PBS). The brain and eyes were removed and post-fixed overnight at 4°C with 4% paraformaldehyde in PBS, followed by more than 24 hours of immersion in 30% sucrose in PBS. Tissues were then embedded in OCT (Tissue Tek, Sakura Finetek). 20 μm sections were cut on a cryostat.

### Immunohistochemistry

Tissue sections were directly placed on positively charged microscope slides (Globe Scientific). Tissue sections were fluorescently immunostained for the following markers: anti-tdtomato: 1∶500 rabbit polyclonal anti-DsRed (Living Colors), immature neuronal marker anti-Doublecortin (1:500, abcam) proliferative marker anti-BrdU (1:200, Abcam). The AlexaFluor tagged secondary antibodies used were Alexa488 (green), Alexa568 (red), and Alexa633 (far red) (Molecular Probes). Antigen retrieval was used for neuronal markers (50 mMol Tris HCl, 45 minutes at 80 °C) and BrdU marker (2N HCl, 0.1% triton, 30 minutes in 37°C)

In preparation for staining, slides with sections were placed in 50 mM Tris HCl for 45 minutes at 80°C. After slides were cooled, slides were incubated in 2N HCl (0.1% Triton) at 37°C for 30 minutes and rinsed in 0.1 M boric acid (pH 8.5) at room temperature for 10 minutes. Sections were then rinsed in PBST (0.2% Triton X-100/ PBS), blocked for one hour with block solution (2% bovine serum in PBS/0.1% Triton X-100), and incubated overnight with anti-BrdU antibody and anti-Hu antibody in serum solution (1% bovine serum in PBS/0.1% Triton X-100) at 4°C. Sections were washed in PBST, incubated with secondary antibodies in serum solution at RT for 2 hours, washed in PBST, counterstained with DAPI, and cover slipped with Vectashield© antifade mounting medium (Vector Laboratories).

### Cell quantification

Tissue sections used for BrdU quantification were imaged using an inverted fluorescent microscope (Olympus IX71). Exposure was kept constant for each channel within experiments. Four to six representative dentate gryi from coronal sections were counted per mouse subject. The number of BrdU+ cells were counted within the dentate gyrus of a 20x image (Figure 3C). BrdU is a maker for proliferating cells. The BrdU+ cell counts were standardized to the volume of dentate gyrus tissue in the image (mm^3^). Volume was determined by measuring the area of the dentate gyrus on ImageJ and multiplying the area by the thickness of the tissue. For analysis of neurogenesis, Doublecortin+BrdU+ and BrdU+ cell within the dentate gyrus were counted. Doublecortin is a marker for immature neurons [45]. The tissue sections used for quantification were imaged using confocal microscopy (SP8 confocal microscope, Leica). To determine the frequency of BrdU+ cells expressing DCX, dual fluorescence-labeled sections were examined by confocal microscopy using a 20x objective (Figure 3F, S6). Sections were scored for single or double labeling by manual examination of optical slices. Cells were manually counted for double labeling when DCX labeling was unambiguously associated with a BrdU+ nucleus. Cells were spot-checked in all three dimensions by Z-stack using a 20x objective. In all cases, the observer was masked to the treatment and genotypes.

### Comprehensive Lab Animal Monitoring System

For analysis of whole animal activity levels and sleep behavior, *Bbs1*^*M390R/M390R*^ mice (n=7) and control mice (n=9) were placed in a Comprehensive Lab Animal Monitoring System (CLAMS; Columbus Instruments, Columbus, OH, USA). CLAMS is an open circuit system that directly measures various parameters over a 72 hour period including movement, sleep behavior, food intake, VO2, VCO2, and heat production. Mice were weighed before the CLAMS recording. Mice were individually housed in Plexiglas cage chambers that were kept at 24°C under a 12:12 hour light-dark cycle. The chamber had 0.6 liters of air passed per minute. Movement (activity) was measured by XY laser beam interruption, and sleep behavior was measured as minimum movement for four minutes or longer. Food consumption was monitored by electronic scales. For measuring the O2 and CO2, the gas content of the exhaust air from each chamber was compared with the gas content of the ambient air sample. The V◻O2 and V◻CO2 measurements were normalized to mouse body weight. The following parameters were calculated as followed: RER = V◻CO2/V◻O2, heat production = 1.232*VCO2+3.815*VO2. CLAMS were performed at the University of Iowa Fraternal Order of Eagles Diabetes Research Center Metabolic Phenotyping Core.

### Slice preparation and electrophysiology

Hippocampal slices from group-housed, naive 2-months old *Bbs1*^*M390R/M390R*^ mice (n=4) and control mice (n=4) were prepared as previously described [87]. First, tissue brain blocks were affixed to the cutting stage, submersed in cutting solution, and transversely sectioned at 400 μm on a Vibratome 1000 Plus (Vibratome, St. Louis, MO). After bisecting into hemispheres, slices were transferred to a holding chamber containing artificial cerebrospinal fluid (aCSF). After 30 minutes, the holding chamber was removed from the water bath and held at RT (22°C) for the remainder of the experiment.

To record field excitatory post-synaptic potentials (fEPSP), aCSF-filled borosilicate electrodes (Corning #0010 glass, resistance <1 MΩ) were positioned in the stratum radiatum of area CA1. Synaptic responses were evoked by stimulation of Schaffer collaterals with bipolar tungsten electrodes (0.1 MΩ, parylene coated; World Precision Instruments, Sarasota, FL). Signals were amplified (AxoClamp 900A Amplifier, Axon Instruments, Foster City, CA), filtered at 1 kHz, digitally-sampled at 10 kHz (Axon Digidata 1440), and stored for offline analysis in Clampfit 10 (Molecular Devices, San Jose, CA).

An input-output curve (initial slope of fEPSP plotted against stimulus intensity) for assessment of basal synaptic transmission was first generated by delivering pulses of 0.2 ms duration every 15 s at increasing stimulation intensities to elicit synaptic responses. Stimulation intensity was then adjusted to yield 40 - 60% of the maximal fEPSP amplitude. The input/output curve is primarily used to calibrate the setting for LTP of the tissue, along with assessing the viability of the tissue.

After acquiring stable baseline responses for 15 minutes, LTP was induced by a theta-burst stimulation protocol consisting of 12 bursts of 4 pulses at 100 Hz. Synaptic responses were sampled every 15 s for 1 h after induction. For analysis, the initial slope of each fEPSP was normalized to the average baseline slope. Time-matched, normalized slopes were then averaged among slices from animals of the same genotype for comparison and plotted as an average of four consecutive responses (*i.e.*, responses sampled over 1 minute). Slices with maximal fEPSPs of less than 0.5 mV, disproportionately large fiber volleys, substantial changes in fiber volley amplitude during LTP recordings, or unstable synaptic responses during baseline or LTP recordings were excluded.

## Statistical Analysis

Statistical analyses were performed using GraphPad Prism 8.0 (GraphPad Software, San Diego, CA). For comparison of two groups, we ran a two-tailed Welch’s t-test. The Welch’s t-test is recommended over the Student t-test because Welch’s t-test performs better when the sample sizes and variances are unequal between groups, and gives similar results when sample sizes and variances are equal [88, 89]. Preliminary tests of equality of variances to determine t-tests are not recommended since it impairs the validity of the Welch’s t-test [90]. For data that appear skewed, we also ran a Mann-Whitney-Wilcoxon Test. For multiple comparisons, we ran multiple t-tests (analyzed each row individually and did not assume consistent standard deviations) corrected for multiple comparisons using the Holm-Sidak method. A two-way ANOVA was used for comparisons of multiple groups with two different independent variables. We then ran a Sidak post-hoc analysis for the relevant data. A three-way ANOVA was used for comparison of multiple groups with three different independent variables. Graphs were generated on GraphPad Prism 8.0. Data are presented as mean with the error bars indicating standard error of means (unless otherwise noted).

## Author Contributions

Conceived and designed the experiments: TP, CSC, RT, AP, JW, VCS, HS, and QZ. Performed the experiments: TP, TV, CSC, CS, YH, CCC, and RG. Analyzed the data: TP, CSC, SCH, NM, CCC, RG, RT, VCS, and KW. Contributed reagents/materials/analysis tools: CSC, SCH, NM, YH, AP, and JW. Wrote the paper: TP, CSC, HS, QZ, RT, JW and VCS.

## Acknowledgements

We thank Valerie Bouffard and Janelle Garrison for the animal husbandry and genotyping. We thank Chantal Allamargot and Kathy Walters, in the Central Microscopy Research Facility at the University of Iowa, for technical assistance in microscopy. We thank Paul Ranum and Richard Smith for their technical assistance with the Auditory Brainstem Response. We want to acknowledge Jamie Soto of the Metabolic Phenotype Core at the University of Iowa, for her technical help with CLAMS. We thank Hamza Farooq for drawing Figure 1a. We thank AJ Chowdhury for assisting with editing of the manuscript. We would also like to acknowledge Denise Harder for her administrative support.

