## Supplemental Figures for "A mouse model of Bardet-Biedl Syndrome has impaired fear memory, which is rescued by lithium treatment"

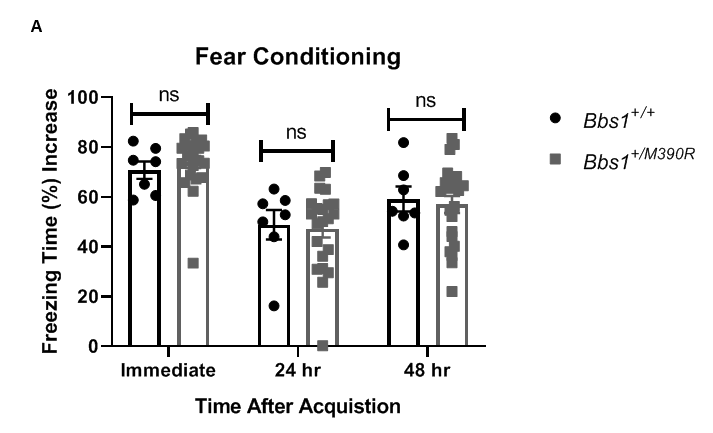

**S1 Fig. Fear conditioning of Wild-type mice (*Bbs1^+/+^*) and Heterozygote mice (*Bbs1^+/M390R^*).** The data was collated from Figure 1 and Figure 6.

The immediate fear conditioning indicates training to the day 1 fear conditioning. The immediate fear conditioning was measured as the freezing time (%) increase of the freezing time (%) just after conditioning (last minute) to the freezing time (%) just before conditioning (first three minutes). The 24 hr fear conditioning represents cue fear conditioning, and was measured as the freezing time (%) increase of the freezing time (%) during the tone (cue, day 2) to the freezing time (%) before the tone (cue, day 2). The 48 hr fear conditioning represents context fear conditioning, and was measured as the freezing time (%) increase of the freezing time (%) during the context on day 3 to the freezing time (%) just before conditioning (first three minutes of day 1).

A.) The immediate fear conditioning was not significantly different between the Wild-type mice (n=7) and Heterozygote mice (n=22) (Welch’s t-test, P=0.494432). The 24 hr fear conditioning (cue) was not significantly different between the Wild-type mice (n=7) and Heterozygote mice (n=22) (Welch’s t-test, P=0.814487). The 48 hr fear conditioning (context) between the Wild-type mice (n=7) and Heterozygote mice (n=22) was not significantly different (Welch’s t-test, P=0.746392).

hr = hours, ns = not significant, * P< 0.05

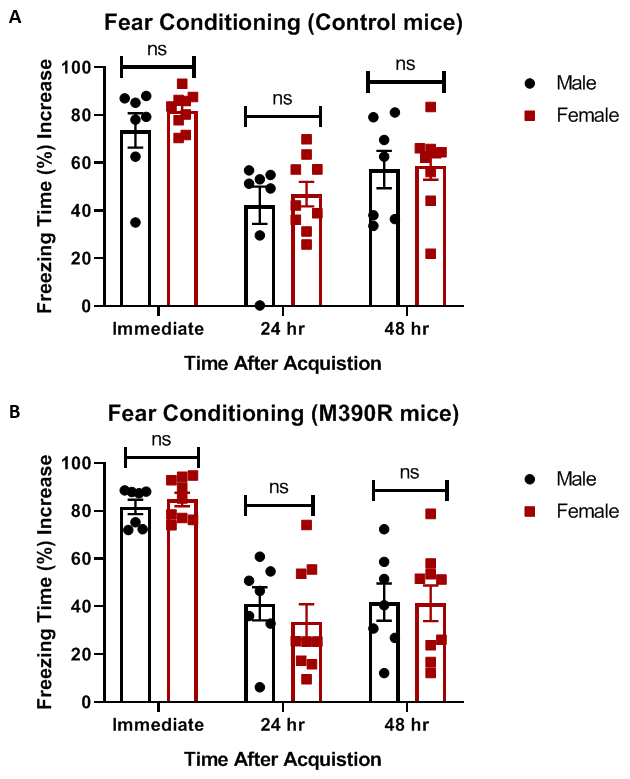

**S2 Fig: No sex effects on fear conditioning in control and *Bbs1^M390R/M390R^* mice.** All data in figure S2 was from the same data pool as Figure 1.

The immediate fear conditioning indicates training to the day 1 fear conditioning. The immediate fear conditioning was measured as the freezing time (%) increase of the freezing time (%) just after conditioning (last minute) to the freezing time (%) just before conditioning (first three minutes). The 24 hr fear conditioning represents cue fear conditioning, and was measured as the freezing time (%) increase of the freezing time (%) during the tone (cue, day 2) to the freezing time (%) before the tone (cue, day 2). The 48 hr fear conditioning represents context fear conditioning, and was measured as the freezing time (%) increase of the freezing time (%) during the context on day 3 to the freezing time (%) just before conditioning (first three minutes of day 1).

A.) The immediate fear conditioning was not significantly different between the Male control mice (n=7) and Female control mice (n=9) (Welch’s t-test, P=0.251805). The 24 hr fear conditioning (cue) was not significantly different between the Male control mice (n=7) and Female control mice (n=9) (Welch’s t-test, P=0.610210). The 48 hr fear conditioning (context) between the Male control mice (n=7) and Female control mice (n=9) was not significantly different (Welch’s t-test, P=0.877612).

B.) The immediate fear conditioning was not significantly different between the Male *Bbs1^M390R/M390R^* mice (n=7) and Female *Bbs1^M390R/M390R^* mice (n=9) (Welch’s t-test, P=0.470202). The 24 hr fear conditioning (cue) was not significantly different between the Male *Bbs1^M390R/M390R^* mice (n=7) and Female *Bbs1^M390R/M390R^* mice (n=9) (Welch’s t-test, P=0.476866). The 48 hr fear conditioning (context) between the Male *Bbs1^M390R/M390R^* mice (n=7) and Female *Bbs1^M390R/M390R^* mice (n=9) was not significantly different (Welch’s t-test, P=0.963440).

control mice= *Bbs1^M390R/+^* mice, M390R mice=*Bbs1^M390R/M390R^* mice, hr = hours, ns = not significant, * P< 0.05

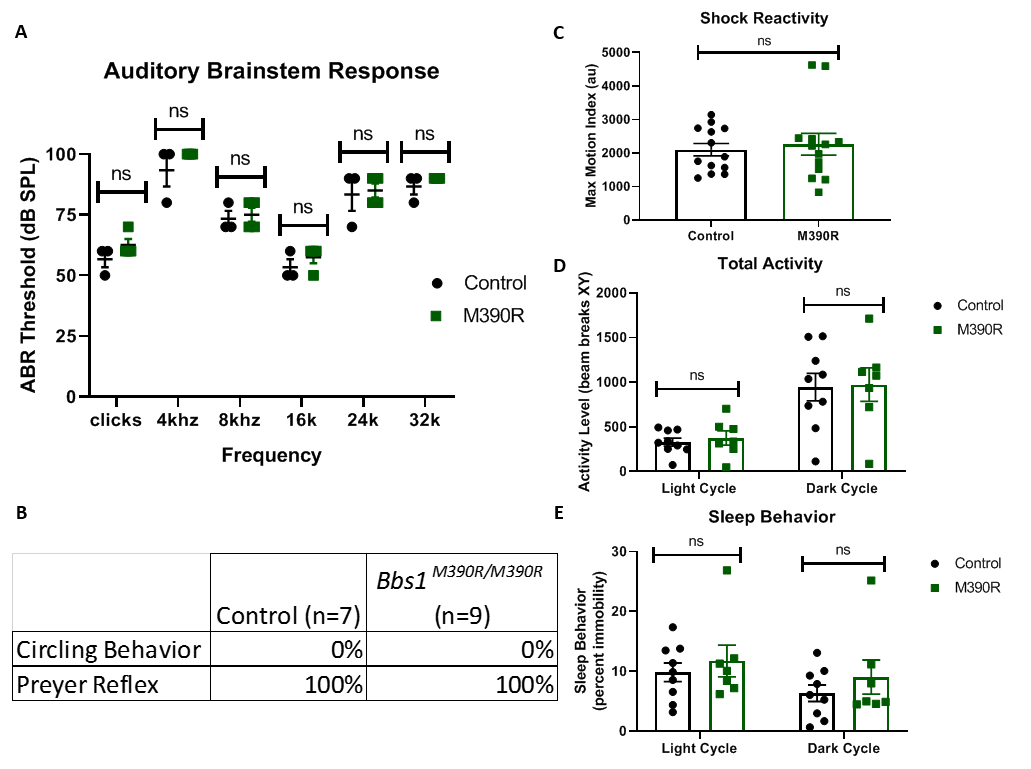

**S3 Fig. Hearing and behavioral assessments of *Bbs1^M390R/M390R^* mice.**

A.) Graph of Auditory Brainstem Response. The threshold of the Auditory Brainstem Response for the control mice (n=3) and *Bbs1^M390R/M390R^* mice (n=4) were not significantly different in clicks (P=0.758), 4khz (P=0.813), 8khz (P=0.922), 16khz (P=0.813), 24khz (P=0.922), and 32khz (P=0.813). Comparisons were analyzed using multiple t-test (without assumption of consist standard deviation), corrected with the Holm-Sidak method.

B.) Table of behavioral hearing test. Both the control mice (n=7, female n=2) and *Bbs1* M390/M390R mice (n=9, female n=4) had intact Preyer reflex and no circling behavior. ns = not significant ABR= Auditory Brainstem Response

C.) Maximum motion index to the first shock in fear conditioning (Shock Reactivity). The control mice (n=13) and the *Bbs1^M390R/M390R^* mice (n=13) were not significantly different in shock reactivity (Welch’s t-test, P=0.672).

D.) The control mice (n=9) and the *Bbs1^M390R/M390R^* mice (n=7) were not significantly different in total activity at Light Cycle (P=0.830) or Dark Cycle (0.908). Comparisons were analyzed using multiple t-test (without assumption of consist standard deviation), corrected with the Holm-Sidak method.

E.) The control mice (n=9) and the *Bbs1^M390R/M390R^* mice (n=7) did were not significantly different for sleep behavior at Light Cycle (P=0.608) or Dark Cycle (P=0.608). ). Comparisons were analyzed using multiple t-test (without assumption of consist standard deviation), corrected with the Holm-Sidak method.

control = *Bbs1^M390R/+^* mice, M390R=*Bbs1^M390R/M390R^* mice, ns = not significant * P< 0.05, ** P< 0.01, ***P<0.001, ****P<0.0001

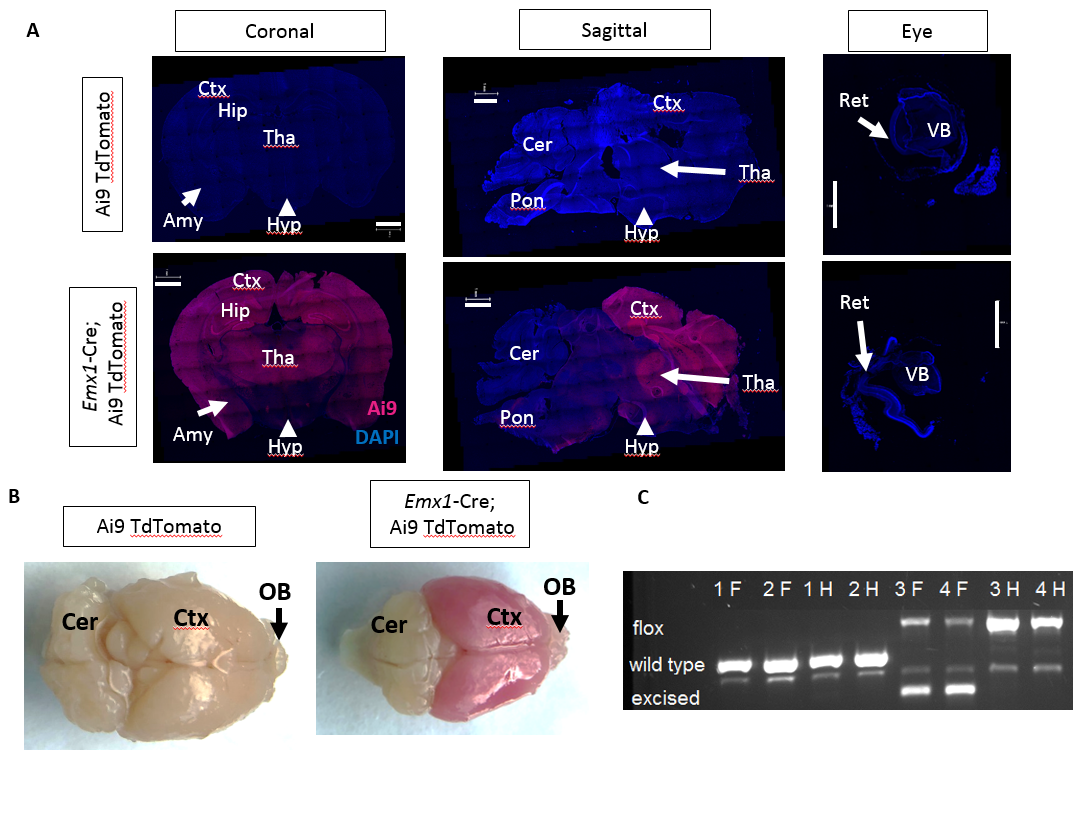

**S4 Fig. *Emx1-Cre* mice.**

A.) Preferential Cre expression in the forebrain. Tissue sections from mice with either Ai9 TdTomato or Ai9TdTomato and *Emx1*-Cre. Ai9 mice do not have any ectopic expression of red fluorescent protein in the brain and retina. Ai9 mice with *Emx1*-Cre have preferential expression of red fluorescent protein in the forebrain, and not in the eye. White line on brain section indicates 0.5mm, and white line on retina indicates 1.0mm

B.) Preferential Cre expression in the forebrain. Brain tissue from mice with either Ai9 TdTomato or Ai9TdTomato and *Emx1*-Cre. Ai9 mice do not have any ectopic expression of red fluorescent protein. Ai9 mice with *Emx1*-Cre have preferential expression of red fluorescent protein in the cortex and olfactory bulb.

C.) DNA gel. Excision band selectively present in the forebrain of *Bbs1*^flox/flox^ ; *Emx1-Cre*+ mice. F=Forebrain, H=Hindbrain, 1, 2 = *Emx1-Cre*+ mice, 3, 4 = *Bbs1*^flox/flox^ ; *Emx1-Cre*+ mice

Cer=Cerebellum, Ctx= Cortex, Hyp=Hypothalamus, Tha=Thalamus, Amy=Amygdala, Hip=Hippocampus, OB=Olfactory Bulb, Ret=Retina, VB=Vitreous Body
